# Reliable reconstruction of cricket song from biophysical models and preserved specimens

**DOI:** 10.1101/2024.10.09.617503

**Authors:** Ryan Weiner, Sarah Duke, Gabriella Simonelli, Nathan W. Bailey, Natasha Mhatre

## Abstract

Predicting the function of a biological structure solely from its morphology can be a very powerful tool in several fields of biology, but especially in evolutionary reconstruction. In the field of invertebrate acoustic communication, reconstructing the acoustic properties of sound-producing forewings in crickets has been based on two very divergent methods, finite element modelling (FEM) and vibrometric measurements from preserved specimens. Both methods, however, make strong simplifying assumptions which have not been tested and the reliability of inferences made from either method remains in question. Here we rigorously test and refine both reconstruction methods using the well-known *Teleogryllus oceanicus* model system and determine the appropriate conditions required to reconstruct the vibroacoustic behaviour of male forewings. We find that when using FEM it is not necessary to assume simplified boundary conditions if the appropriate parameters are found. When using preserved specimens, we find that the sample needs to be rehydrated for reliable reconstruction, however, it may be possible to accomplish rehydration *in silico* using FEM. Our findings provide a refined methodology for the reliable reconstruction of cricket songs, whether from fossils or preserved specimens from museums or field collections.

## Introduction

Animal acoustic communication signals are extensively studied due to their important role in mediating social interactions such as cooperation, conflict, learning, parental care, sexual selection, and reproductive isolation [1,2]. Understanding the causes of acoustic signal variation can be assisted by structure-based reconstruction: in cases where taxa are extinct or unavailable, but archival or fossil specimens remain, reliably reconstructing acoustic signal properties is critical for accurate inferences about signal development, function, and evolution. In most vertebrates, vocalisations are based on soft vocal-tract tissues which are under significant neuro-muscular control [3]. In contrast, invertebrate acoustic communication typically relies on vibrations produced by hardened, cuticular structures. For example, crickets produce loud mate attraction calls using specialised hardened forewings [4]; in addition to mate attraction, these calls and their reception by conspecifics function in mate recognition, courtship, sexual selection, and aggression [5]. The structural and mechanical properties of hardened cricket wings control most aspects of call structure, such as frequency, duration and even loudness [4,6,7]. Thus developmental or evolutionary processes that alter wing structure can drive signal evolution, and hard tissues if preserved can carry a record of signal evolution amenable to reconstruction [8,9]. Refining methods of structure-based signal reconstruction from preserved specimens would therefore accelerate our understanding of signal evolution in cricket species [10–12].

An area of particular interest is the effect of wing venation patterns on cricket song frequency. Wing veins are used to delineate the harp and mirrors which act as resonators in song production [13]. A modified vein, the file, is part of stridulatory apparatus that excites these resonances using a highly regulated clockwork mechanism [7]. There is now considerable evidence that changes in wing venation patterns are genetically driven, and that these changes have been a strong driver of observable signal evolution in crickets [10–12,14]. Some researchers have argued that vibrational patterns from preserved specimens can be used to infer the patterns of vibrations in live crickets because venation is the structural feature that is critical [15], however, there is no direct empirical or modelling evidence showing that this is the case.

Most biomechanical models which predict cricket wing resonances from structure typically use the finite-element (FE) method [9,16–18] and make strong simplifying assumptions about how veins provide the boundary conditions of the system. Current models assume that all the external boundaries of the resonating wing are clamped or held immobile. They justify this assumption by arguing that in regions of the wing with a high density of veins, we typically observe high stiffness and low movement levels in vibrometric measurements and it is therefore reasonable to consider these boundaries to be immobile or ‘clamped’ [16,17]. However, in real wings, only the base of the wing is truly clamped and other wing edges are free to move [13]. It has not been tested whether FE models with the correct boundary conditions capture the reduction in vibration observed high vein density areas. Indeed, we do not have a systematic way of deciding at what vein density would a part of a wing become immobile and for what frequency range.

For instance, in *Teleogryllus oceanicus* from Hawaiian populations, a mutation called *flatwing* silences males and is maintained by selection pressure from the acoustically-orienting parasitoid fly *Ormia ochracea* [14]. Silent *flatwing* males have partially-feminised forewings with very high density of veins throughout the wing and a very reduced harp [10]. Therefore, if we used the boundary condition proposed by existing FE models, we would have to assume that all regions of the *flatwing* wing are stiffened and immobile, a circular statement without any need for an FE model. Additionally, this model class would have little to say about the newer wing variants observed in *T. oceanicus* which have a range of vein densities and wing sizes [14,19]. Indeed, some of these wing variants have reduced but functional singing structures like the file and plectrum [19], and others possess vestigial movements associated for singing [20] which may give rise to novel forms of sound production which would be missed by current methods. Thus, we cannot use the simplifying assumptions currently used in finite element models of cricket wings for realistic structural reconstruction of wing resonances across the substantial diversity of forewing venation patterns and densities found both within and across cricket species. To accurately reconstruct the transition from non-singing to singing species, there is a considerable need for models that explicitly model these stiffening structures and their effects of wing resonances and vibration.

While it might be the case that stiffening structures such as veins determine the spatial pattern cricket wing vibrations, it is not clear that they determine the frequency of this vibration as has been assumed in studies of preserved specimens, another method used for the reconstruction of acoustic function [15]. It is well known that insect cuticle becomes stiffer when desiccated, as captured by an increase in the Young’s modulus, and also that it experiences reduced damping [21,22]. Increases in wing cuticle stiffness will also increase the stiffening effects of the veins since they are made of the same material. Thus, resonances measured from preserved specimens are expected to be at higher frequencies than those from live animals. Indeed, resonance frequencies cannot be predicted from morphological features alone; the Young’s modulus of insect cuticle varies considerably [23], and so do simple morphological features like wing cuticle thickness, both of which affect resonance frequency [18].

To fill this gap in our understanding of how cricket wing resonances are determined by their structure, we developed a finite element model of *Teleogryllus oceanicus* wings where we used realistic boundary conditions, clamping only the wing base and explicitly modelling the stiffening effect of wing veins using a coupled plate and rod formulation. We began by developing a model of wildtype (‘normal-wing’, Nw) males using known morphology and measurements of wing vein diameters (Fig. S1). We then used a process of fitting and iterative refinement to infer two unknown mechanical properties which are responsible for the resonance frequency, i.e. the Young’s modulus of wing cuticle (*E*) and the thickness of the wing membrane (*mt*) (Fig. 1b). Next, we used further refinement to estimate the damping in the system. We used these mechanical properties and the known morphology of *flatwing* (Fw) mutant wings to test if a model of these wings predicts loss of resonance observed in the real insects [10].

**Figure 1:**
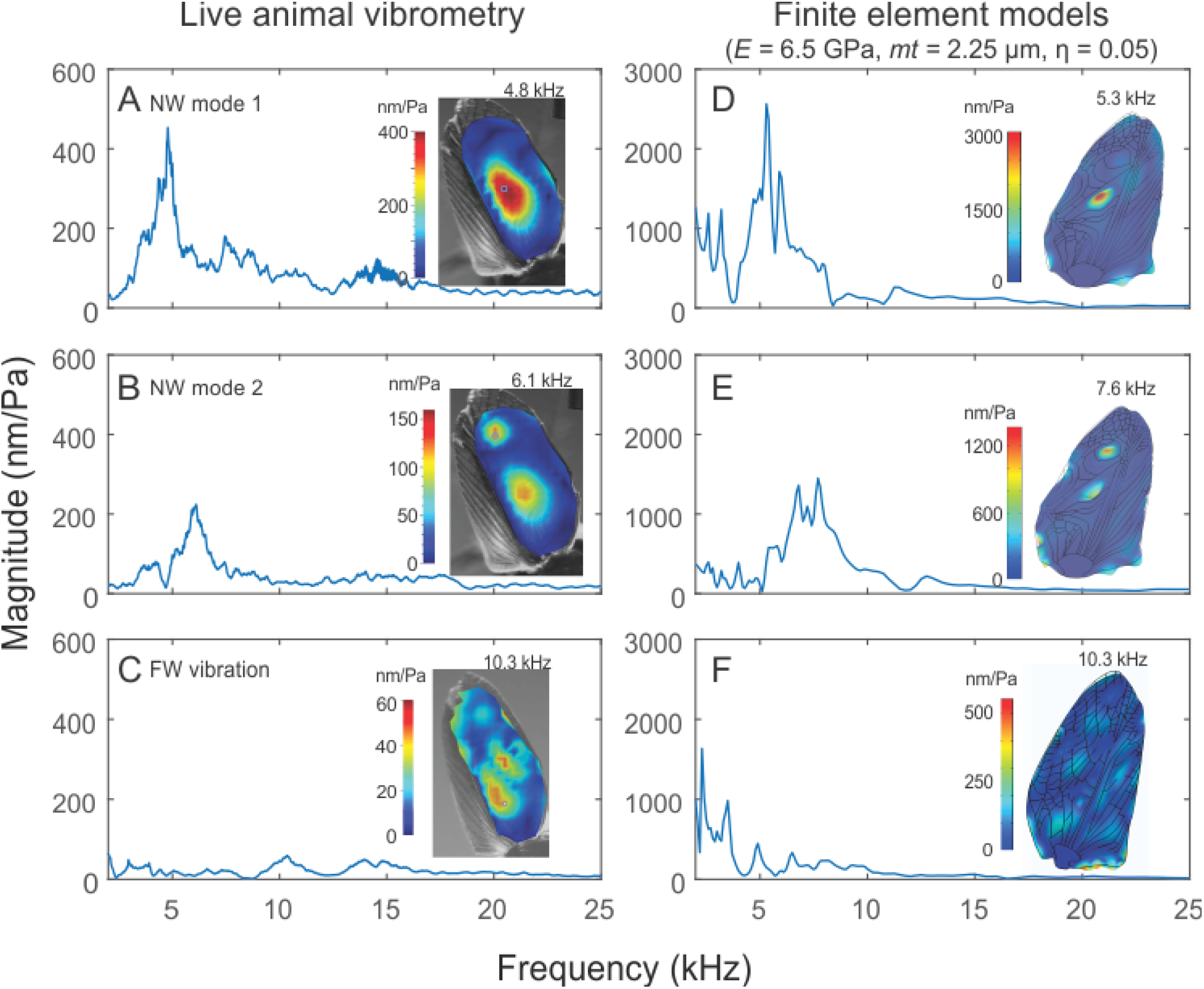
Finite element models capture the vibrational patterns of real *Teleogryllus oceanicus* wings. The singing forewings of *wildtype* or normal winged *T. oceanicus* crickets have (A) have a sharp peak in their frequency response near their singing frequency which corresponds to a resonant mode which we refer to as harp mode or mode 1. To observe this peak in the frequency response, we plot the frequency response at a central point on the harp which is depicted in the figure. (B) Their forewings have another resonant mode at a higher frequency where the harp vibrates out of phase with the mirror cells. To observe the appropriate frequency response for this mode, we plot the frequency response of a central point on the mirrors which is depicted in the image. (C) In *T. oceanicus* crickets that carry one of the *flatwing* mutations, the forewing has an altered venation structure, an altered frequency response with significantly lower vibration amplitudes. A finite element model which incorporated the appropriate geometry of the normal *T. oceanicus* wing and had a realistic parameter set for the wing cuticle modulus and thickness showed (D) a harp mode at a song like frequency, and had a very similar frequency response from the same point on the harp in terms of both the resonant frequency and the Q factor (see results). The same model also reproduced (E) mode 2 in which the harp and mirror show anti-phase vibrations. The frequency response of the modeled mirror resembled that of the real mirror in terms of both the resonant frequency and the Q factor (see results). When the same model parameters were applied to the wings of flatwing mutants, the model predicted vibration shapes that were very similar to those observed in real wing and which had very similar low amplitude frequency responses.

Additionally, we conducted a second test of our models and test whether we can also predict the resonance behaviour of dry preserved wing specimens and tested their relationship to live specimens. Dry preserved insect cuticle is known to become stiffer, with an elevated Young’s modulus, and experience lowered damping [21,22,24]. These parameters can be independently modified in our model and would allow us to estimate accurate values for the resonances of preserved wing specimens and their relationship to live wings. Thus, using a combination of finite element modelling and vibrometry, we can identify which features of the singing structures of cricket wings can be reliably inferred from preserved or fossil specimens for the reconstruction of signals and evolutionary analysis.

## Methods

### Finite element model construction

Both FE models were developed in COMSOL Multiphysics (v5.5a, Burlington, MA, USA). Detailed methods are provided in the supplementary materials. Vector drawings of *T. oceanicus* wings and venation patterns were imported into COMSOL to provide model geometry. The wing veins were defined as beams completely coupled along their length to the wing membrane which was defined as a plate with defined thickness. Vein thicknesses were measured using optical coherence tomography (OCT) (Fig. 1 A; see supplementary materials for detailed methods). Wing membrane thickness was below OCT resolution, and was found through iterative refinement, along with the Young’s modulus of the wing and vein cuticle. For damping we initiated the model with Rayleigh damping parameters derived from vibrometry measurements as described before [16] and then iteratively refined damping estimates using the isotropic loss factor formulation. Typically, insect wing veins are a mix of solid structural veins and hollow veins that carry blood and the axons from sensory neurons distributed across the wing [25,26]. In the models reported here all veins were treated as solid rods, since this replicated real behaviour. When the veins were treated as hollow pipes with a wall thickness set at the same level as membrane thickness, the models did not replicate wing behaviour suggesting that in cricket wings modified for singing, veins are predominantly solid and structural in nature.

First, we conducted a series of eigenfrequency studies in COMSOL for all parameter combinations under consideration (Fig. S3). An eigenfrequency study finds the resonance frequencies of the modelled system and the corresponding spatial vibrational patterns or modes at each eigenfrequency. Then we identified the subset of these parameter combinations that were close to model fit criteria. Model criteria were based on the mechanics of wildtype *T. oceanicus* wings and included 1) the prediction of a resonant mode in which only the harp vibrated between 4.5-55 kHz with a Q factor of 5, and 2) a second mode between 5.8-7.7 kHz where the mirrors also showed significant anti-phase vibrations, which had a magnitude at least 1.7 times lower than the first resonant mode (see supplementary methods for details). Only resonance frequencies and mode shapes can be identified exactly in eigenfrequency studies. Therefore only these part of the criteria could be evaluated in these studies. For the subset of parameter combinations that partially met model criteria, we conducted frequency domain studies which are more computationally expensive but generate estimates of the Q factors and also the relative amplitudes of different modes. In these frequency domain studies we examined the vibrational behaviour of the wings in response to a constant pressure (20 mPa, or 60 dB SPL re. 2e-5 N/m^2^) applied to the wing across a range of frequencies (2-25 kHz). Outputs from this study were then used to examine the frequency response of different points on the wing (Fig. 2).

**Figure 2:**
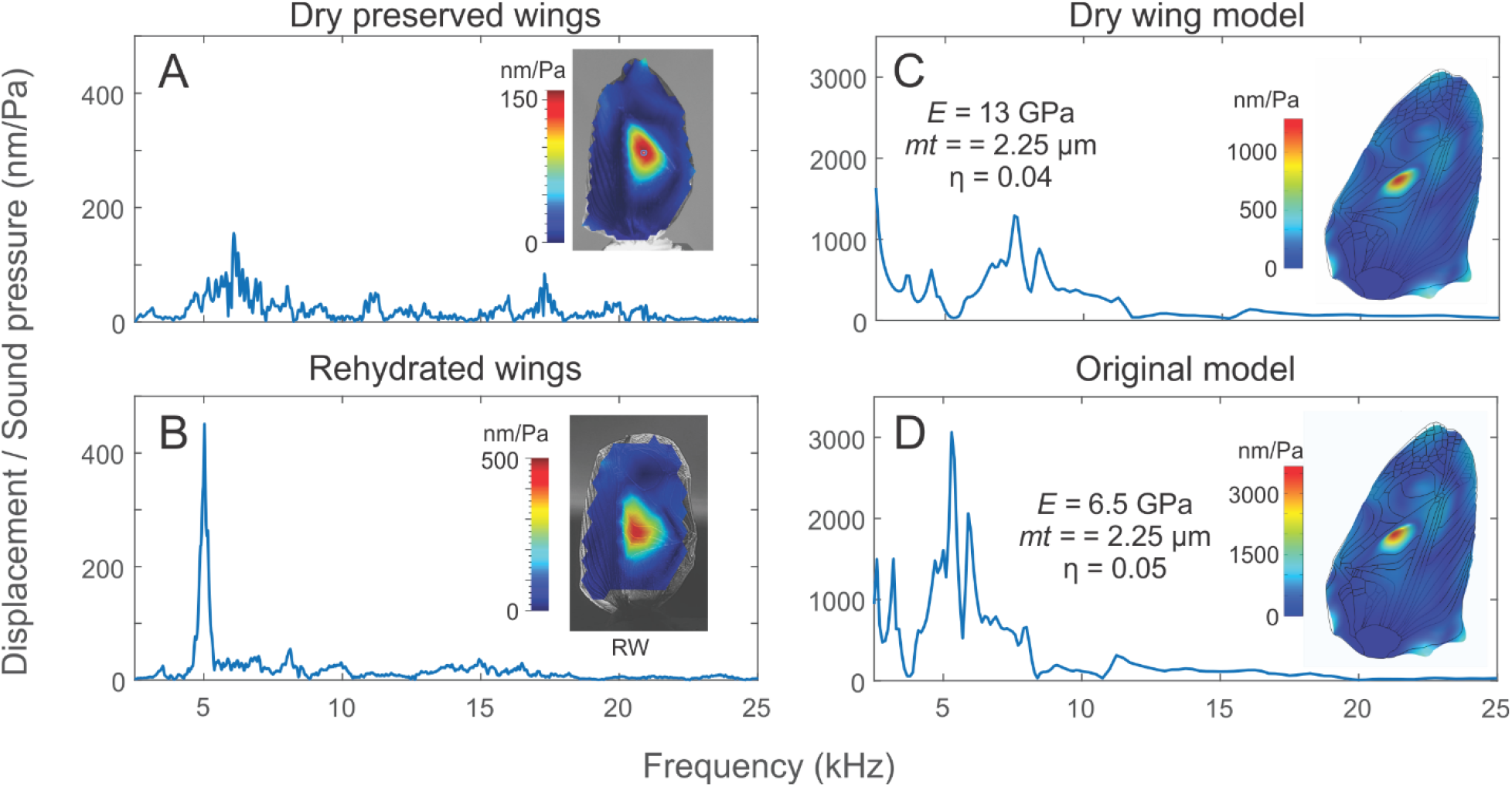
Dry preserved forewing specimens can be informative about the vibrational mechanics of live animals. (A) LDV measurements from dehydrated dry-preserved *T. oceanicus* wings have a harp mode. However, this mode occurs at a higher frequency than ones observed in live animals as can be observed from the frequency response at the harp. (B) When rehydrated, the harp mode is retained. and the resonance frequency decreases as can be observed from the frequency response at the harp, and is the same as that observed from live animals. (C) A finite element model with a Young’s modulus that is twice that of the (D) live normal wing model captures the frequency response observed at the harp of dehydrated dry-preserved wing, coupled with a small decrease in damping. This is consistent with previous observations that the Young’s modulus of desiccated insect cuticle is known to increase and its damping is known to decrease [21,22,24]. Thus, both data and models suggest that models and dry-preserved wing specimens could be used to reconstruct the vibrational mechanics of cricket forewings following careful analyses.

### Vibrometry of preserved wing specimens

In 2019, forewings were collected from normal-wing and flatwing males obtained from laboratory *Teleogryllus oceanicus* lines. The lines were pure-breeding for the respective genetic variants and were originally derived from a population in Kauai, Hawaii where the two morphotypes segregated at the time of collection [see [27]. After removal with dissecting scissors, forewings were mounted on cardboard with a small piece of Blu Tack (Bostik), placed in an insulated tube, and maintained dry until use approximately 5 years later. Dry preserved wings were rehydrated as needed by submerging into distilled water for about 1 hour directly before measurement. To facilitate comparisons, we used laser Doppler vibrometry to measure the vibrational behaviour of both dry and rehydrated forewings in response to acoustic stimulation at a range of frequencies similar to those used for live animals [10]. These data were used to identify the harp mode, and its modal frequency, amplitude, and the Q-factor of the modal frequency response (see supplementary materials for detailed methods). All three metrics were compared between dry and rehydrated wings using paired t-tests in R (RStudio Version 1.4.1717). Similarly, all three metrics data were also compared with vibrometry data from live *T. oceanicus* wings which were published in a previous study [10] using unpaired t-tests. Data from the forewings of live animals were used as a baseline to evaluate the effects of preservation and subsequent rehydration.

## Results

### Models of normal wings

We used data from vibrometric measurements of the wings of living wildtype individuals from a previous publication [10] to assess whether the model output fit real wing behavior (see supplementary methods for details). In the first iteration of eigenfrequency studies for the normal-wing or Nw model (Fig. 1 B), we tested a broad range of parameters (Young’s modulus *(E)* = 1-10 GPa and membrane thickness (*mt*) = 2-10 μm) at a low resolution (1 GPa and 1μm, respectively). These model parameter ranges were based on previously reported data from other cricket species [18]. The model was first set to output eigenfrequencies between 4 and 6 kHz and the resulting mode shapes for each parameter combination.

We examined these eigenfrequencies to see if they possessed a mode which was isolated to the harp with an anti-node centred in the harp which fell within the natural range, i.e. between 4.5-5.5 kHz.

Next, we ran a second iteration of eigenfrequency studies (Fig. 1B) using a narrower range of parameters (*E* = 5-10 GPa and *mt* = 1-5 μm) surrounding the few that fulfilled the first criteria. These were run at the same resolution as the previous iteration (1 GPa and 1 μm, respectively) however, the eigenfrequency search was now extended and went from 4 to 10 kHz. Within this range, we checked whether the parameter combinations also produced a second mode with antiphase movement in the mirrors. We found that the *E* values which would reproduce the main harp modes between 4.5-5.5 kHz and showed some movement in the mirrors at higher frequencies were between *E* = 6 and 7 GPa and when membrane thickness was either 2 or below 3 μm. Eigenfrequencies do not give us a clear indication of the full frequency response and therefore of the Q factor of the resonances. They also do not predict which modes are superposed on each other when driven at a particular frequency. Therefore, further parameter refinement was achieved through a frequency domain study.

To identify which parameter combination best captures Nw wing resonance, we conducted a frequency domain study searching for *E* = 6-7 GPa and *mt* = 2-2.5 μm, at a higher resolution of 0.5 GPa and 0.25 μm, respectively (Fig. 1B). The frequency responses for a point on the harp and for a point on the mirror-cells for each combination of *E* and mt were analyzed to see which parameter combination best met our criteria. The first part of our criteria was that the harp’s frequency response should show a single peak frequency between 4.5-5.5 kHz (5.00 ± 0.52 kHz, mean ± SD, n=4, n = 4 for 2 real animals, 2 wings) with a Q-factor of approximately 5.0 (5.51 ± 0.68, mean ± SD, n=4, n = 4 for 2 real animals, 2 wings). We also set the criteria that the mirror mode or the second mode in which the mirrors move in an antiphase fashion to the main harp will occur at a frequency ∼1.3 times higher, and between 5.8-7.7 kHz (6.58 ± 0.90, peak frequency ratio: 1.3±0.05, n = 4 for 2 real animals, 2 wings) than the harp but at least 1.7 times lower in magnitude (harp mode: 438.34± 63.77 nm/Pa, mean ± SD; mirror mode: 138.93± 86.58 nm/Pa, mean ± SD; peak height ratio: 4.88 ± 3.91, mean ± SD, range: 1.7 to 10.1, n=4 for 2 animals, 2 wings). The mirror mode is highly variable in real animals and is not always the peak frequency of the mirrors, hence we consider this a weak criterion.

Additionally, we do not expect exact matches in displacement levels at either mode from the models and do not use these in our model fit criteria. In the vibrometry measurements, the wings are stimulated by a sound which will applies force on both faces on the wing suspended in the sound field, therefore the exact net force level on the wings is unknown. In the model, we apply pressure only in one direction and the force levels and therefore the absolute displacements between the model and real wings cannot be directly compared. We do, however, expect that the relative displacements between the harp and the mirror mode will be similar between the model and the real wings.

The best combination of parameters was found to be *E* = 6.5 GPa and *mt* = 2.25 μm (Fig. 2A,B) which are congruent with data from other cricket species [18]. Initial values of damping used were then updated and we found that an isotropic loss factor (η) of 0.05 gave us the best fit between models and real animals. At the harp, the highest peak frequency occurred at 5.3 kHz with a magnitude of 2562 nm/Pa. The shape of the vibrational mode predicted by the model was similar to that seen in the real data, with a single high amplitude anti-node present in the centre of the harp with smaller associated movement in the mirrors.

The Q-factor of this vibrational mode was ∼5.88, similar to that seen in real measurements. At the mirror-cells, there was a peak frequency centered around 7.6 kHz with a magnitude 1097 nm/Pa. This is at a frequency that is 1.4 times the harp resonance frequency and the magnitude of the mirror resonance is 2.3 times lower that the harp’s resonance. The shape of this vibrational mode predicted by the model was also similar to that seen in the real data, with two out of phase anti-nodes in the harp and the mirror respectively. Thus, in our model the parameter combination of *E* = 6.5 GPa, *mt* = 2.25 μm and an isotropic loss factor (η) of 0.05 provided the best prediction of the resonance of *T. oceanicus* normal wings in terms of capturing wing resonant frequencies and modal shapes.

### Models of *flatwing* mutants

Using this parameter combination, we modelled the flatwing *T. oceanicus* mutant. The frequency response for a point at the vestigial harp in the Fw rod model displays numerous very small peak frequencies and a large overall reduction in movement that is comparable to the observed reduction in forewing vibration amplitude in comparison ot Nw males (Fig.1) as reported in a previous study [10]. In the real data we see some displacement localised to the vestigial harp at 10.3 kHz, with similar movement levels at other positions on the wing. The model also shows similar movement at the vestigial harp and across the wing at 10.3 kHz. Thus, our Fw FE model also matches the mechanical behavior of the real

Fw, indicating that venation pattern is primarily responsible for determining the spatial pattern of wing resonance in *T. oceanicus* wings.

### Measurements and models of preserved wing specimens

We then measured the vibrational behaviour of *dehydrated* dry-preserved specimens of Nw *T. oceanicus* collected from the Kauai lineage that had been dry preserved since 2019. We found that that when excited by sound, these wings had a harp mode that appeared very similar to that of live Nw *T. oceanicus* wings (Fig. 3A). The harp mode in these desiccated wings occurred at a significantly higher frequency (6.76 kHz ± 0.84, N=10) than that observed in live specimens (5.0 kHz ± 0.53, N=4, unpaired t-test, t-stat=-4.24, P = 0.0006). It is well known that the Young’s modulus of insect cuticle increases when excised and dehydrated [21,22,24], thus this is not an unexpected result. We additionally observed a decline in displacement levels in dry preserved wings (230.68 ± 52.91 nm/Pa, N=10) in comparison with live wings (438.34± 52.91 nm/Pa, N=4, unpaired t-test, t-stat=-7.05, P =0.0001) consistent with an increase in stiffness. The wing modes also had a higher Q factor in dry preserved wings (51.04± 24.63, N=10) than live wings (5.51± 0.68, N=4, unpaired t-test, t-stat=4.24, P=0.001) suggesting an expected decline in material damping, however, this was harder to estimate due to the ‘rough’ nature of the frequency response. These data are consistent with a model with Young’s modulus that has increased by a factor of 2 to 13 GPa, and a lower isotropic loss factor of ∼0.04.

**Figure 3.**
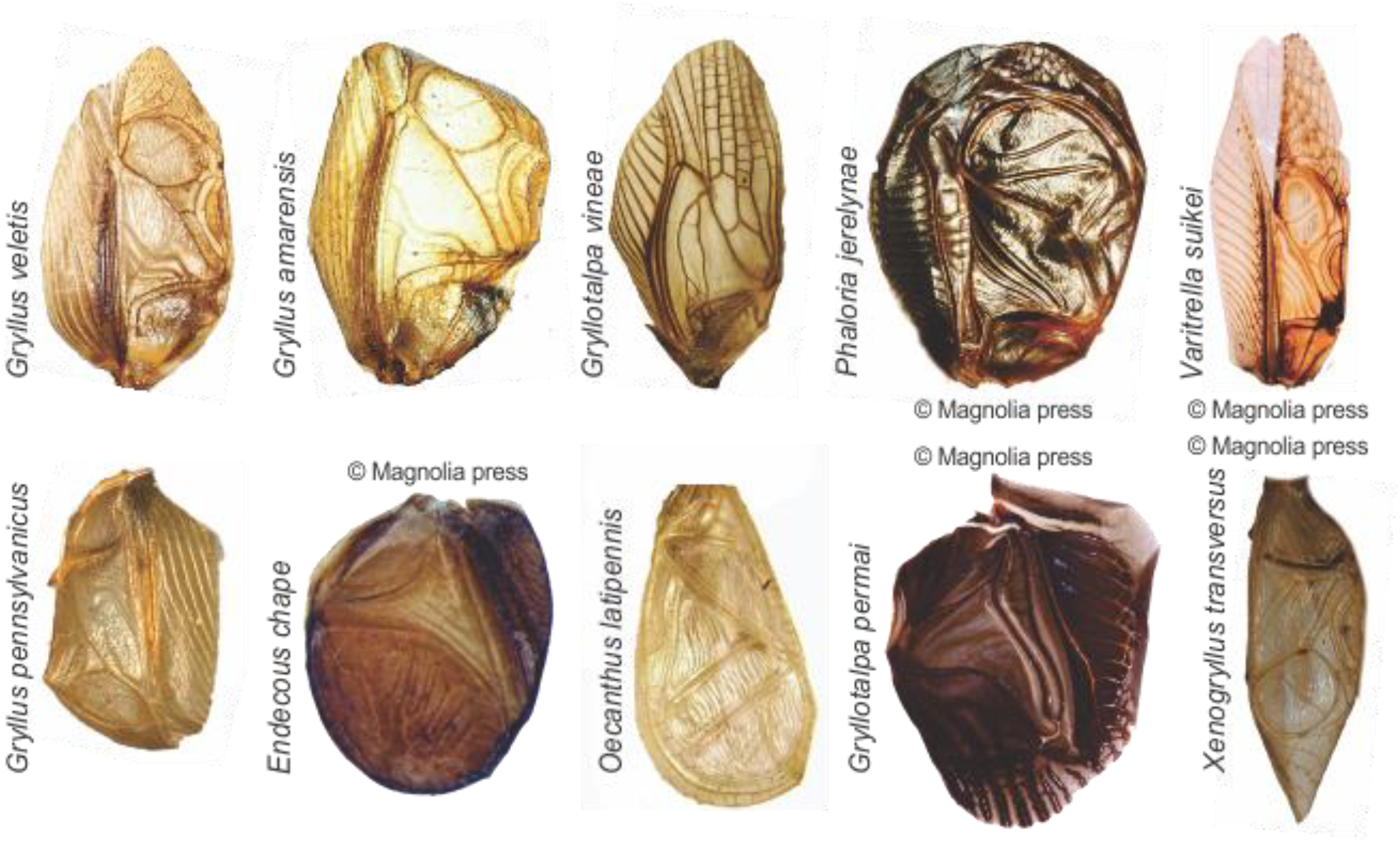
A few example cricket forewings from acoustically active members of the true crickets (Grylloidea) depicting the diversity of wing shape, venation and sclerotization observed across the phylogeny. Permissions to reproduce images: *Gryllotalpa vinae*: Museum D’Historie Naturelle ID: MNHN-EO-ENSIF4425 (CCBY: RECOLNAT (ANR-11-INBS-0004) - Marion DEPRAETERE - 2017); *Gryllus veletis*: Orthoptera Species File Collection object 1499333 (CCBY-SA: © 2009 University of Michigan of Zoology, Holger Braun); *Gryllus pennsylvanicus*: (Specimen SEM-UBC GRY-0542, Photo by Don Griffiths reproduced courtesy of the Spencer Entomological Collection, Beaty Biodiversity Museum, UBC); *Gryllus amarensis*: Museum D’Historie Naturelle ID: MNHN-EO-ENSIF7031 (CCBY: MNHN - Ranjana JAISWARA - 2018); *Oecanthus latipennis:* University of Guelph Insect Collection: Specimen BIOUG44550-E07 (CCBY: CBG photography group, Centre for Biodiversity Genomics); *Endecous chape:* (reproduced with permission from the copyright holder, Fig. 2A: Souza-Dias et al., 2017: https://doi.org/10.11646/zootaxa.4237.3.2); *Phaloria jerelynae*: (reproduced with permission from the copyright holder, Fig. 2: Gorochov and Tan, 2012: https://doi.org/10.11646/zootaxa.3525.1.2); *Gryllotalpa permai:* (reproduced with permission from the copyright holder, Fig. 2E: Tan and Kamaruddin, 2016: https://doi.org/10.11646/zootaxa.4066.5.3); *Varitrella suikei*: (reproduced with permission from the copyright holder, Fig. 16D: Tan et al, 2020: https://doi.org/10.11646/zootaxa.4810.2.2); *Xenogryllus transversus*: (reproduced with permission from the copyright holder, Fig. 5G: Jaiswara et al., 2019: https://doi.org/10.11646/zootaxa.4545.3.1)

The Young’s modulus of dessicated insect cuticle is also known to return to live values upon rehydration [21,22]. When rehydrated for a period of ∼1 hour, the frequency of the harp mode (Fig. 2B) lowered to frequencies (5.08 ± 0.25 kHz, N=10) similar to those of live crickets (5.00± 0.53 kHz, N=4, unpaired t-test, t-stat=-0.28, P = 0.79). The displacement levels of the rehydrated wings remained lower and were more variable (175.12 ± 133 nm/Pa, N=10, unpaired t-test, t-stat=-5.79, P = 8.77e-5) and the mean Q factor also remained elevated (16.55 ± 7.78, N=10, unpaired t-test, t-stat= 4.12, P =0.001), suggesting that material damping remains high even after rehydration.

## Discussion

### Venation & spatial pattern of vibration

Our data show that FE models based on the rod formulation can capture the shapes of cricket forewing resonances, without additional constraints or strong boundary condition assumptions. Additionally, comparisons between the vibration patterns observed in *T. oceanicus* normal wings and *flatwing* mutant models where no other parameter was changed, supports the idea that high vein densities reduce vibration locally and suppress the formation of modes, except a small antinode observable in the harp (Fig. 1). Thus our models show that while it is not possible to *a priori* predict the vein density required to significantly suppress local vibration, it is possible to predict vibration patterns by explicitly modelling real veins with realistic boundary conditions.

One relevant limitation of our analyses is that we assumed that all veins are either rod- or pipe-like in nature, whereas in reality a wing may have some combination of both. We observe in our model that the harp mode is somewhat isolated to the higher half of the harp whereas, in the real wing, there is considerable vibration over the cross veins in the harp as well (Fig. 1, 2). This suggests that these veins in the interior of the harp are less stiff than the veins along the boundary. We also make the assumption that the wing is of uniform thickness, whereas a local thinning of the cuticle may also contribute to movement within the harp. Nonetheless, our data suggest that wing venation and its stiffening effects are critical to forming the appropriate spatial patterns of resonant excitation, known as resonant modes. The harp mode is critical because the harp is bordered by the file vein and this vein must vibrate in order to enable the clockwork mechanism which couples and maintains the plectrum drive at the resonant frequency [4,7,13]. Thus making a reliable prediction of the shape of the harp mode or any mode that would excite the file vein is important since it predicts which frequencies could shape the clockwork mechanism. For instance, both harp modes and higher frequency vibrational modes of forewings might be important to the production of calling song as has been speculated in some Eneopterines [28] and observed in tree crickets [16]. In tree crickets, the spatial pattern of these higher modes can be seen to involve the file vein, explaining the ability of the tree cricket clockwork mechanism to work at a broader frequency range [16].

In *T. oceanicus*, both the harp mode and the mirror modal shapes remained similar across a wide range of cuticle stiffness as predicted by the model and demonstrated by the dry-preserved specimens (Fig. 2). This suggests that variations in stiffness caused either by cuticle sclerotization during development or due to hydration status would not significantly alter the spatial pattern of vibrations across individual crickets. It also suggests that spatial patterns of vibration are possible to infer from desiccated specimens, at least where these modal patterns are the lower frequency high-amplitude modes of the forewings such as the harp. We found that the harp mode of *T. oceanicus* wings was present in both stiffer models and dry-preserved forewings, whereas the mirror modes, which are at higher frequencies were more variable in the dry-preserved forewings. Thus, where higher frequency vibrational modes of forewings might be important to the production of calling song as has been speculated in some Eneopterines [28], the use of dry-preserved or even rehydrated specimens may not be ideal.

### Reliability of inferences from preserved specimens

Vibrometry measurements from dry preserved specimens of *T. oceanicus* wings, which are expected to have increased cuticle stiffness due to dessication, were indeed found to have increased resonant frequencies but retained a harp mode (Fig. 2). This is congruent with our models which also predict that increases in wing cuticle modulus cause increases in resonant frequency but maintain the harp model shape (Fig 2). The frequency responses measured from both the model and the dry preserved specimens were significantly noisier than from fresh wings, which is likely caused by the lower damping observed in dehydrated insect cuticle [21,22]. Thus not the just the frequency but also the shape of the ‘resonance filter’ of cricket wings is altered by dry preservation. Thus, while it is possible to make some inferences about the spatial patterns of vibration in a dry preserved cricket wing specimen, it is not possible to infer the shape or the frequency of the resonant filter underlying the cricket’s song.

Interestingly, however, we found that rehydration lowered the resonant frequency of dry-preserved wings so that it was the same as that observed in live animals (Fig. 2). Previous work has shown that the increased Young’s modulus in dehydrated wings, can be restored to its normal value simply by rehydration [21,22]. Indeed, our data shows that rehydrated wings have vibrational mechanics that are statistically indistinguishable from those measured from live animals, where the harp mode is concerned.

This raises the interesting possibility that dry-preserved specimens may be used for investigating wing mechanics, with the caveat that the effect of dehydration and rehydration should be tested for a wider range of cricket wing morphologies (Fig. 3). A range of possibilities open up if this proves to be the case, such as systematic studies of cricket song biomechanics without the need for live specimens. As an example, it would be possible to examine the mechanics of dry preserved museum specimens, where paratypes are available to be manipulated via rehydration rather than risking critical type specimens. It would also be possible to study the wing mechanics of new species as they are found without the need for live specimen transfers. Such mechanical studies would further taxonomic characterization in addition to song and could be used to track natural variation and incipient speciation. An additional possibility is to examine mutational variations observed in natural populations[29] or induced in experimental genetic studies, also without the need for shipping live specimens across national boundaries. Finally, it would also be possible to model the mechanics of the wings of fossil crickets enriching existing approaches [8].

### Modelling cricket wing diversity and song evolution

It is worth reiterating at this point that there is a huge diversity of cricket wing structures across the true cricket phylogeny (Fig. 3). However, even among the Gryllines, the relative shapes and sizes of the harp and the mirrors can vary greatly as can be observed in the three members of this genus *Gryllus, G. veletis, G. pennsylvanicus* and *G. amarensis* (Fig. 3). Indeed across the Gryllid phylogeny, members of which all sing in a similar fashion to Gryllines, we see a huge variation in parameters which are likely to affect wing mechanics: wing size, shape, sclerotization, and wing membrane thickness (Fig. 3)[6,18,30]. In particular, there is a huge variation in the venation pattern that different species use to stiffen their wings and therefore to determine wing resonance. In contrast, the diversity of true cricket (Gryllidae) wing mechanics that have been studied so far is relatively small, there are some measurements of wing mechanics of Gryllines [10,13,31], of Oecanthines [16] and of some Eneopterines [32].

Our study which explores how venation pattern determines vibration mechanics and cricket song therefore opens up the possibility of a model based approach to studying cricket song diversity. One can quite easily imagine the use of transformation grids, such as those used by D’Arcy W. Thompson [33], or more modern landmarking techniques [34,35] to describe the variations observed in cricket wing venation diversity across the phylogeny in existing databases [6]. Such transformation grids could be used to develop FE models. Existing information about the song frequency of different species [6] could then be used to tune models allowing us to infer Young’s modulus & cuticle thickness for a range of species. The predictions of selected models could be tested, especially those of unusual morphologies or very distinct wing types such as those of Oecanthines. These measurements could be made with live and with dry preserved specimens, to further test the utility of our findings here. The results of such a large scale analysis would enable us to disentangle the interactions between the main morphological determinants of song frequency in the gryllid phylogeny, whether development has favoured wing modulus, size, shape, or membrane thickness as the main lever for signal divergence.

## Acknowledgements

We would like to thank the following funding sources for their support of this research: Canadian Foundation for Innovation (CFI4748) and the Ontario Research Fund, NSERC Discovery (Grant no. 687216), and early career supplement (675248), and an NSERC Canada research chair (Grant no. 693206) to NM; the Western SEED grant to NM; Western Undergraduate Work Study to SD; Western Undergrad Summer Research Internship to RW; and UK Natural Environment Research Council funding to NWB (NE/I027800/1, NE/L011255/1, NE/W001519/1).

## Notes

### Competing Interest Statement

The authors have declared no competing interest.

### Summary of Updates

Figure 3 has been updated. We have now obtained permissions from the copyright holders of all copyrighted images and have updated the copyright statement as appropriate. No other changes were made.

