## Supplementary materials for "Reliable reconstruction of cricket song from biophysical models and preserved specimens" (from another version of this preprint: version 2, 2024-10-13)

### MATERIALS AND METHODS: FINITE ELEMENT MODEL CONSTRUCTION

Measuring vein thickness:

Images of *T. oceanicus* wings were imported into an annotation program, Notability (Version 11.7.2, Ginger Labs Inc., San Francisco, California, USA). We then developed a vein labelling system consistent with the Comstock-Needham system [1]. Veins not captured in this system were also annotated and names were based on wing region (Fig. S1).

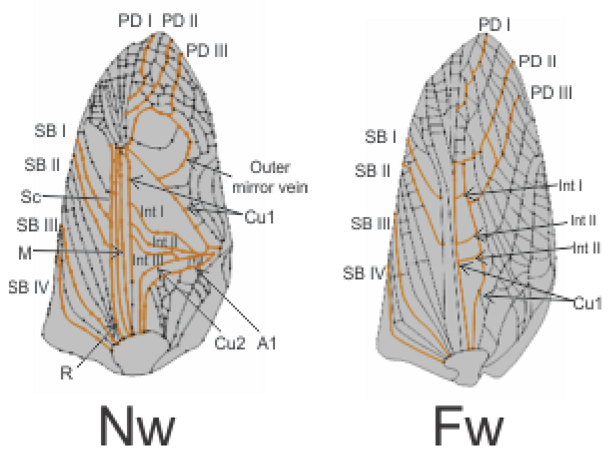


Figure S1: Major vein names in the forewings of the wildtype normal wing (Nw) veins and the equivalent veins in the flatwing (Fw) mutant forewing.

The vein thicknesses of 5 pairs of wings from Nw Kauai males, and Fw Kauai males were measured using optical coherence tomography (OCT). OCT is a non-invasive optical imaging technology that can produce cross-sectional images of an object (Podoleanu, 2012). Using the Ganymede 620F OCT system (GAN620C, ThorLabs Inc., Germany) sections of each individual vein were scanned at a resolution of 2 μm. We then scanned each labelled vein three times, with each scan being spaced out at three different locations along the vein. This allowed us to calculate average vein thickness to account for variation along the entire vein. Recorded OCT scans were exported as .TIFF files and processed in an image processing software (Fig. S2).

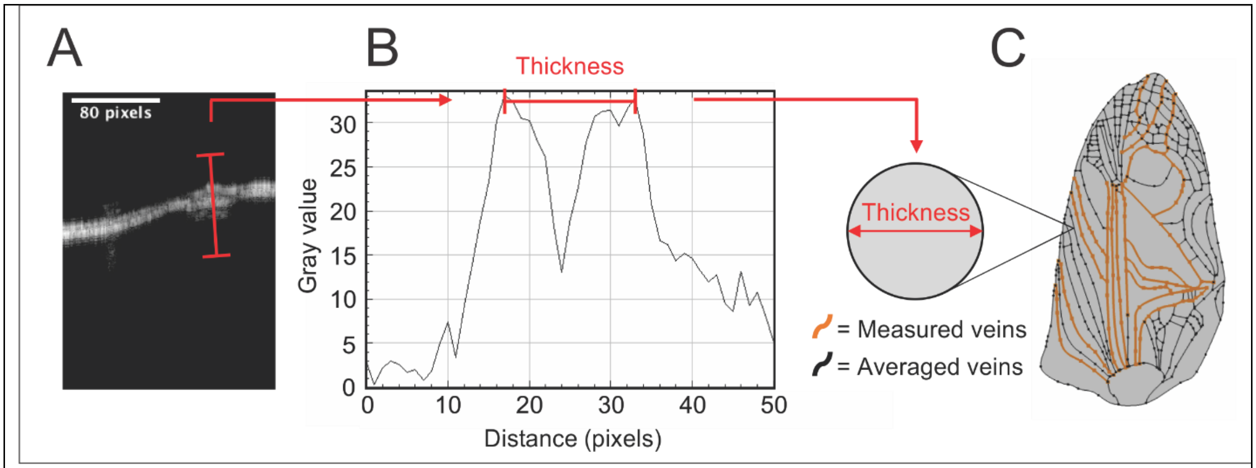


Figure S2: Vein thicknesses were measured using optical coherence tomography images.

Calculating average vein thickness:

ImageJ (Version 1.53k, National Institutes of Health, USA), a Java-based imaging processing program, was used to make measurements of vein thickness. OCT uses a broadband laser and measures the light levels reflected by biological tissue, through its depth. Denser tissue reflects more light, and air does not reflect any light allowing us to differentiate between tissues of different density but also tissue and air. In our case only the wing cuticle reflected any light. Using these data, we could make measurements of vein thickness using a gray-scale plot of reflectivity along the beam depth and sweep (Fig S2 A). The distance between the first two grayscale peaks along beam depth were measured using imageJ and taken to be the vein thickness in micrometers (μm) (Fig S2, Table S1).

Table S1: The average vein thickness observed in dry-preserved specimens and used for FE modelling

| Normal wing | | | Flat wing | | |
|---|---|---|---|---|---|
| Vein thickness (μm) | | | Vein thickness (μm) | | |
| Vein Name | Average | Standard error | Vein name | Average | Standard error |
| PDI | 28.17 | 4.33 | PDI | 25.17 | 2.69 |
| PDII | 29.25 | 4.31 | PDII | 24.92 | 2.89 |
| PDIII | 28.36 | 4.73 | PDIII | 26.46 | 3.33 |
| SBI | 41.85 | 11.50 | SBI | 26.12 | 6.17 |
| SBII | 40.22 | 11.67 | SBII | 26.73 | 4.30 |

| | | | | | |
|---|---|---|---|---|---|
| SBIII | 39.49 | 4.01 | SBIII | 35.44 | 6.73 |
| SBIV | 37.74 | 3.51 | SBIV | 33.39 | 6.00 |
| Outer Mirror Vein | 42.56 | 10.10 | Cu1 | 35.75 | 4.76 |
| Cu1 | 48.01 | 7.47 | | | |
| IntI | 41.58 | 8.43 | | | |
| IntII | 42.38 | 11.03 | | | |
| IntIII | 40.13 | 10.69 | | | |
| Cu2 + File | 45.26 | 6.16 | | | |
| A1 | 47.08 | 7.56 | | | |
| R | 39.29 | 18.55 | | | |
| M | 45.37 | 22.43 | | | |
| Sc | 38.48 | 17.00 | | | |

Finite element modelling:

All finite element modelling in this manuscript was carried out using COMSOL Multiphysics (version 5.5 and 6.1, COMSOL Incorporated, Burlington, MA, USA). Finite element modelling is often used to model the mechanical behaviour of structures with a complex geometry. In general, finite element modelling is an approximation method that takes a complex structure and subdivides it into a finite number of smaller “elements”, whose mechanics are described by a set of partial differential equations [2].

In the first stage we define the problem to solved, i.e. the mechanical object that we wish to model. Problem definition includes defining the geometry of the object, the properties of its materials, the boundary conditions, and the forcing conditions it experiences. Next we must choose the solver module which will solve the problem and this determines the equations which will be used which we will predict its behaviour. To model the normal wildtype *T. oceanicus* forewing (Nw), we started by defining the geometry, i.e. the shape of the wing, and the positions and thicknesses of the veins using data in the previous section ((Figure S1, Table S1). The thickness of the stiff membrane (*mt*) between the veins was treated as a free parameter and varied across models. Then we defined the material properties of these different parts, including the material’s density ($\rho$=1200 kg/m$^3$ based on previous estimates of cuticle density [3]), Poisson’s ratio ($\upsilon$=0.3, the ratio observed in most materials [4]), and Young’s modulus ($E$) was a free parameter and varied across models based on previously reported values in cricket forewing cuticle [5,6]. The damping experienced by the wing was then set using Rayleigh damping ratios calculated from previous vibrational analysis of *T. oceanicus* wings (f1 = 4825, $\zeta_1$ = 0.08, f2 = 33883, $\zeta_2$ = 0.18) as reported before [4]. In subsequent iterations, damping was updated using the isotropic loss factor formulation in as needed to obtain a better fit. Next, the wing's boundary conditions were set to be "clamped" only at the wing-base, and all other boundaries were free in order to simulate the wing being attached only to the cricket's body. Depending on the type of study the forcing conditions were also defined and are described later. The final step in this stage was to generate the mesh which divided the wing into a series of interconnected finite elements. The wing mesh was set to physics controlled and of normal size, i.e. it had a maximum element size of 1 mm and a minimum element size of 0.18 mm. The final mesh contained 5664 triangular elements with an average quality of 0.74.

The inputs provided in the first stage define the mechanics of the wing, which are then described by a set of mathematical model equations defined by the software. COMSOL uses a formulation known as the mixed interpolation of tensorial components (MITC) in modelling the wing membrane of the 'shell' domain and the veins are modelled using the Euler Bernoulli formulation and the two are coupled using a shell-beam connection and solves for displacement of wing and vein elements as the final output.

Model analysis and fit criteria:

To quantify model fit, we developed criteria which would indicate whether the FE models accurately represented the vibrational behaviour of real *T. oceanicus* forewings. Measurements of real wing vibrations obtained using Laser-Doppler vibrometry (LDV) from a previous study were used [7]. Specifically, we had data from four wings from two Nw individuals and eight wings from four Fw individuals.

To assess the vibrational behaviour of forewing, we extracted the frequency responses of a central point on the harp and a central point on the mirror-cells in the right and left wings of both Nw males (*N* = 2 individuals, 2 wings) using PSV software (Polytec software (v. 10.1.1.3, Polytec GmbH, Waldbronn, Germany)). In all individuals and all wings, we identified the wing resonance by identifying the peak in the frequency response of the harp (peak frequency). Each peak frequency fell within the range of known Nw songs (4.5-5.5 kHz). We also calculated the Q-factor of the resonance of each wing. Q-factors are a unitless parameter describing the sharpness of a resonance. They were calculated by hand by dividing the frequency at which the peak occurs ($f_{peak}$) by the difference in the frequencies at half the peak's magnitude ($\Delta f$) (as seen in Equation 1).

$$Q = \frac{f_{peak}}{\Delta f} \quad (1)$$

Where the available transfer function did not contain the exact value of half the peak's magnitude, the closest value near the peak was taken. The higher the Q-factor, the sharper the peak. The average Q-factor at the point on the Nw harp was approximately 5.51 ± 0.68 (n = 4 for 2 real animals, 2 wings). We used these two features to set the first criteria for a good Nw model fit.

***To be considered a good model, the model should predict a peak frequency between 4.5-5.5 kHz, with a Q-factor of approximately 5.0 as measured at a central point on the harp.***

In contrast to the point at the harp, when the transfer function at a central point on the mirror is observed there are multiple peak frequencies, and these peaks differ substantially across individuals and wings. These peaks indicate the mirror mode which can be found among one of these peaks and is the second mode of the wing in which the mirrors move in an antiphase fashion to the main harp. The common feature about the mirror peak frequencies was that they were higher frequencies (ranging from 5.8 to 7.7 kHz, 6.58 ± 0.90, n = 4 for 2 real animals, 2 wings) and the peak had a magnitude lower (1.7 to 10 times, 4.79 ± 3.89, n = 4 for 2 real animals, 2 wings) than that of the harp's peak frequency. Given the high variability of this mode even in wildtype wings, we consider the second Nw model criteria based on mirror-cell behaviour to be less stringent.

***To be considered a good model, the model should reproduce a peak frequency at the mirror between 5.8 and 7.7 kHz with a peak magnitude that is at least 1.7 lower than the harp peak.***

At the harp's peak frequency and at the mirror-cells' peak frequencies, there is movement isolated to the harp and the mirror-cells (Fig. 1A,B), where only these structures move at the frequency of interest and other parts of the wing remain immobile. This modal shape along with the peak in the frequency response indicates that these frequencies are resonant frequencies, which are known to produce the calling song [1,4,7,8]. Therefore, we also looked for this spatial pattern of vibration in the NW model at the harp's peak frequency and at the mirror-cells' peak frequencies

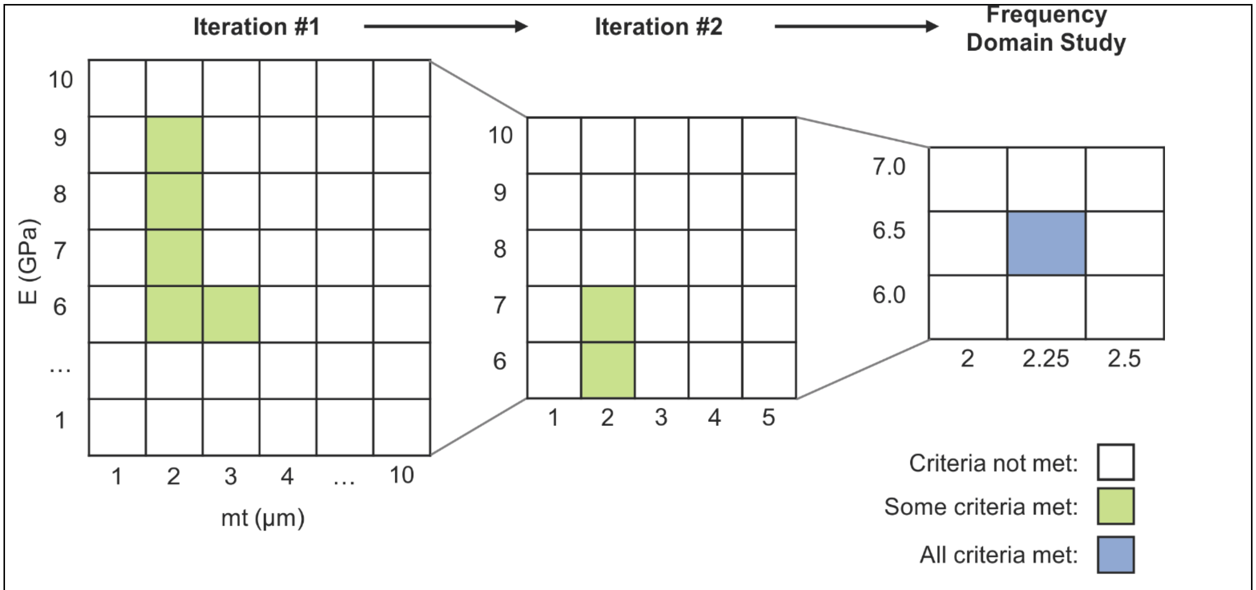


S3: The iterative refinement paradigm used to identify the parameter combination that produced the best model fit.

Construction of the Nw and Fw wing geometry:

Vector drawings of Nw and Fw were developed in a vector graphics editing software CorelDRAW (CorelDRAW X7, Version 17.1.0.572, Corel Corporation, Ottawa, ON, Canada). Using the poly-line tool, wing boundaries and wing veins were traced and represented as a solid line. Wing-bases represent regions were drawn and these will be the area of the wing that is represented as "clamped" to the cricket body, meaning that they will be held stationary in the models.

Construction of the Nw models: Wing and vein geometries:

The updated Nw schematic was exported as a .DXF file and imported into COMSOL Multiphysics software (Version 5.5a or 6.1, COMSOL Inc., Burlington, MA, USA) as a three-dimensional object. To model the forewing, I used COMSOL's shell module where the stiff membrane of the wing is modelled as a linear elastic isotropic plate using the Mixed Interpolation of Tensorial Components (MITC) formulation. The veins are modelled as solid rods using the beam interface using an Euler-Bernoulli formulation. Finally the veins and the shell are connected using the "shell-beam connection" and set as having shared edges along the shell normal, which clamps the modelled beam and the shell together, with the veins stiffening the plate locally. The veins were given diameters based on measurements and remaining veins were given a diameter that was an average of the diameters of neighbouring measured veins (Fig. S1, Table S1).

Nw model parameters:

The material properties used in the NW models must represent plausible values. In past studies, the material density (ρ) and Poisson's ratio (υ) of insect cuticle has been reported to fall in a very small range, we used these values from previous literature [3–6]. The density was defined to be 1200 $kg/m^3$ and the Poisson's ratio to be 0.3. The two parameters that remain unknown for *T. oceanicus* wings are its Young's modulus (*E*), which represents the stiffness of the material, and the thickness of the wing membrane (*mt*). A common method for estimating these values is to sweep through a range of plausible values and refine the estimated value by iterating through different combinations of parameters until the parameter combination that best captures the vibrational behaviour of the target structure is found.

Once a parameter set which reproduced Nw wing vibrational behaviour was developed, we used that parameter set to generate a Fw model. To then determine whether this model reasonably reproduced Fw vibration behaviour we developed a set of criteria to assess Fw model fit. First, we extracted the frequency responses of the vestigial harp in the right wing of 4 individuals ($N = 4$ individuals). This yielded frequency responses appropriate to the mutant wing phenotype (Fig. 1C). The frequency responses for all individuals had multiple peak frequencies below 16.0 kHz, which is consistent with the maximum resonance frequency range in Fw males of 16.0 kHz[17]. Furthermore, the magnitude of all these peak frequencies are approximately 7 times lower than the magnitude at the harp's peak frequency in Nw males, thereby indicating reduced resonant behaviour. At all these peak frequencies, there is modally patterned movement across the entire wing. ***So the Fw model was thought to have met the fit criteria if it had multiple small peaks in its frequency response, and a highly reduced amplitude throughout its frequency response.***….

Nw model parameter selection:

In *T. oceanicus* wings, the values of the wing material's Young's modulus (*E*), an intrinsic property that represents the wing's stiffness, and its membrane thickness (*mt*) are unknown. To find these values, we conducted two different computational studies in the Nw models: eigenfrequency studies and frequency domain studies.

An eigenfrequency study finds the resonance frequencies of the system and plots the corresponding eigenmodes at each of these frequencies. A "mode" is the spatial pattern of the wing at a particular frequency and an eigenmode is a mode at a resonant or 'eigenfrequency'. For example, in Nw males there is a peak frequency between 4.5 and 5.5 kHz. The corresponding mode at this frequency has high movement primarily at the harp, with the rest of the wing remaining immobile.

In contrast, a frequency domain study examines the vibrational behaviour of the wing across a range of frequencies. This is done by applying a constant force to the wing at those frequencies and measuring the wing's displacement at each frequency. The model outputs from a frequency domain study can then be used to produce a frequency response at any point on the wing, which enables us to calculate its Q-factor as well as to quantitate the vibration amplitude. Additionally, we can visualize the spatial pattern of vibration at any of the modelled frequencies and the output will contain any superposition of eigenmodes that would normally occur at frequency.

Eigenfrequency studies take a fraction of the time to run in comparison to frequency domain studies. Therefore, we first ran eigenfrequency studies to search for the best combination of $E$ and $mt$, and then used frequency domain studies with the best combination of parameters to confirm if the models were successful in replicating real Nw wing resonance.

In the Nw model, we initially ran eigenfrequency studies over a large range of combinations of Young's modulus values ($E$ = 1.0–10.0 GPa) and membrane thicknesses ($mt$ = 2.0-10.0 μm), with parameter values spaced far apart at a low resolution (1.0 GPa and 1.0 μm, respectively). The initial range was chosen based on values determined in a previous study[18] and from literature[12]. The model was first set to output only those eigenfrequencies that lay between 4 and 6 kHz and the resulting mode shapes for each parameter combination. After analyzing the mode outputs of each combination of Young's modulus and membrane thickness, testing for criterion 1 (a mode with movement confined to the harp between 4.5-5.5 kHz), we determined which parameter combinations best met these criteria.

As this allowed us to refine our search, we ran a second iteration of an eigenfrequency study. We increased the upper end of the search range of the eigenfrequency study to 10 kHz, and this time also evaluated models for criterion 2 (movement confined to the mirror-cells at higher frequencies) and determined which parameter combinations best met both criteria. The parameter range of the final iteration was $E$ = 6.0-7.0 GPa and $mt$ = 2.0-2.5 µm at a resolution of 0.5 GPa and 0.25 µm, respectively.

We then ran frequency domain studies for these combinations of parameter values. From the resulting model outputs, we produced frequency responses at a central point on the harp and a central point on the mirror-cells. The locations of these points were the same as the points on the harp and mirror-cells used in the analysis of the LDV data. The frequency responses showed the displacement magnitude predicted by the model at each point over a span of frequencies ranging between 2.0-25.0 kHz (at 100 Hz resolution between 2 and 10 kHz and 250 Hz resolution between 10 and 25 kHz). In frequency domain studies, the model needs to be driven by an external force. The model was excited by a uniform face load of 0.02 Pa at all frequencies, which approximates 60 dB SPL (re. 2e-5 N/$m^2$) of sound pressure being applied to only one face of the wing. Thehese data could then be compared to the frequency responses of the real wings obtained using LDV, thereby allowing us to test against the Nw criteria established in the previous section.

Construction and analysis of the Fw model:

The Fw schematic was exported as a .DXF file and imported into COMSOL Multiphysics software. The Fw model was constructed in a similar way to the Nw model and vein dimensions were based on data measured from Fw forewings. The Young's modulus and membrane thickness of the Fw model were taken from the combination of $E$ and $mt$ in the selected Nw model.

A frequency domain study was conducted using the same range of frequencies as used in the NW models (2.0-25.0 kHz). Using the model output, we produced a frequency response for a point on the vestigial harp that was located in the same position as the point on the vestigial harp used in the analysis of the LDV data. These data were then used to test whether the Fw model met the Fw criteria as defined above.

### MATERIALS AND METHODS: PRESERVED WING SPECIMEN ANALYSIS

Forewing origins, preservation and rehydration:

Laboratory stocks of *Teleogryllus oceanicus* from a population in Kauai were established as previously described in detail by Bailey et al., 2019. Select individuals had their forewings removed, mounted on cardboard, placed in an insulated tube, and were preserved for 5 years. The same 5 pairs of Nw male forewings were measured first in the dehydrated, dry-preserved condition and then after rehydration. Wings were rehydrated by submerging each pair of wings in distilled water for 1 hour directly before measurement.

Laser-Doppler Vibrometry:

We measured the vibration of these *T. oceanicus* forewings using Laser-Doppler vibrometer (LDV) (Polytec PSV-500, Polytec GmbH, Waldbronn, Germany). The wings were mounted at the end of a rectangular base attached to the end of a long cylindrical holder and positioned perpendicular to the lens of the LDV laser head. A speaker (FF125WK, FOSTEX, Tokyo, Japan) was placed under the mounted wings to deliver an acoustic stimulus to vibrate the wings in a method similar to that used by previous studies [4,7]. The acoustic stimulus consisted of periodic chirps (frequencies ranging from 2.5 to 25 kHz, at a frequency resolution of 20 Hz) generated using Polytec software (v. 10.1.1.3, Polytec GmbH, Waldbronn, Germany), passed to an amplifier (PM6007, Marantz, Kew Gardens, USA) and sent to the loudspeaker. The amplitude of sound at all frequencies within this range was equalized to 60 ± 0.5 dB (sound pressure level relative to 20 μPa). We positioned a high-precision microphone (Type 2670, Brüel & Kjær, Nærum, Denmark) directly beside the mounted wings to enable equalization and to monitor and record the sound stimulus during the measurement as a reference. The mounted wings were scanned over a grid of 400-500 scanning points, averaging 20 times at each point. Surface vibration measurements were made in velocity per unit of time from which displacement was calculated. Then using a transfer function, the wing displacement with respect to the reference sound pressure was calculated. To monitor measurement noise, the coherence between the vibrometer and microphone signals for each scanning point was also calculated.

Data was analyzed using Polytec software. The aim was to identify the presence of the harp mode, and its vibration characteristics which include the resonant frequency, the amplitude of the resonant peak, and the Q-factor. All three characteristics were measured by hand using Polytec software within the resolution of the measured frequency response. Paired t-tests were performed, to compare all characteristics between the dehydrated dry-preserved and rehydrated specimens. As well, unpaired t-tests were performed to compare both the dehydrated dry-preserved and rehydrated-preserved specimens to the harp modes of live Kauai *T. oceanicus* wings, measured and reported in a previous publication [7]. It should be noted that the sample size of the live specimens (n=4) was smaller than that of the dry-preserved and rehydrated-preserved sample size (n=10).
